# Coordinated social interactions are supported by integrated neural representations

**DOI:** 10.1101/2024.01.16.575885

**Authors:** Silvia Formica, Marcel Brass

## Abstract

Joint actions are defined as coordinated interactions of two or more agents towards a shared goal, often requiring different and complementary individual contributions. However, how humans can successfully act together without the interfering effects of observing incongruent movements is still largely unknown. It has been proposed that interpersonal predictive processes are at play to allow the formation of a Dyadic Motor Plan, encompassing both agents’ shares. Yet, direct empirical support for such an integrated motor plan is still limited. In this study, we aimed at testing the properties of these anticipated representations. We collected EEG data while human participants (N = 36; 27 females) drew shapes simultaneously to a virtual partner, in two social contexts: either they had to synchronize and act jointly, or they performed the movements alongside, but independently. We adopted a multivariate approach to show that the social context influenced how the upcoming action of the partner is anticipated during the interval preceding the movement. We found evidence that acting jointly induces an encoding of the partner’s action that is strongly intertwined with the participant’s action, supporting the hypothesis of an integrative motor plan in joint but not in parallel actions.

## Introduction

A huge variety of human activities can only be accomplished through coordinated interactions with others. Think of two tango dancers: to perform smoothly and elegantly, they need to coordinate their actions in time and space, constantly adjusting their movements to match their partners. Such concerted activities are often referred to as Joint Actions (Sebanz et al., 2006), situations in which two or more agents interact in order to achieve a shared goal, as, for example, performing a *pirouette*. However, the cognitive mechanisms allowing for such coordination are far from being fully understood (Sacheli et al., 2018).

One crucial issue in Joint Actions is that of visuomotor interference (*VMI*, Blakemore & Frith, 2005; Kilner et al., 2003). Observing another agent performing a movement is known to activate the corresponding motor plan in the observer (Rizzolatti et al., 1996), influencing their movement execution. An incongruent observed action produces a disruption of the observer’s motor plan and, consequently, poorer motor performance (Brass et al., 2001; Cracco et al., 2015; Cracco & Brass, 2019; Forbes & Hamilton, 2017). Notably, acting jointly towards a shared goal often requires the coordination of complementary and incongruent movements (Sartori & Betti, 2015). It has been proposed that interference is overcome by generating a Dyadic Motor Plan that integrates the anticipated behavior of the other into one’s own motor plan (Sacheli et al., 2018). Indirect evidence for such an integration process comes from behavioral data on reduced VMI when acting jointly compared to acting alongside but independently (i.e., in parallel), quantified in faster reaction times and lower distortions in the kinematic profiles of the executed movements (Clarke et al., 2019; Rocca et al., 2023; Sacheli et al., 2018, 2019).

Neurocognitive studies addressing interpersonal coordination mostly focused on how acting jointly modulates motor activation following an observed movement, thus inferring predictive processes during action unfolding (Bolt & Loehr, 2021). Only few EEG studies investigated the interval preceding the movements, reporting univariate differences between interactive and non-interactive social contexts (Kourtis et al., 2013, 2014), and highlighting the benefits of anticipating the full joint configuration (Kourtis et al., 2019). Although these studies provide some evidence for differences in neural activity associated with preparing for Joint Actions, they do not test directly the proposed integrated representation of the upcoming movements, and their results might be reflecting more general features of the task, such as difficulty or conflict anticipation.

To fill this gap, we developed a novel Paired Drawing Task, during which we asked participants to draw the instructed shapes either synchronously with a virtual partner (i.e., Joint social context) or in parallel. By controlling visual, motor, and attentional demands of the task, we ensured differences in how the action plans are represented during the preparation interval could only be attributed to their relevance for the upcoming Joint or Parallel action.

We set out to address two main questions. First, we tested whether drawing jointly reduced VMI, as reported in previous behavioral studies (Sacheli et al., 2018). Second, we hypothesized that acting jointly relies on dedicated neural representations, proactively anticipating and tying together each agent’s contribution. Therefore, to directly tap into the representational content of the neural activity recorded during the preparation interval, we implemented a multivariate approach to investigate what information was decodable from the spatial and temporal activation patterns. We reasoned that, if during Joint Actions the upcoming movements of the participant and the partner are represented intertwined, incongruent shapes combinations should be difficult to distinguish from each other, indicating the emergence of an integrative representation as proposed by the Dyadic Motor Plan framework.

## Materials and Methods

### Participants

Forty participants were initially recruited, and gave their informed consent prior to the beginning of the experiment in accordance to the approved application to the Ethical Committee of Humboldt University (reference number: 2022-45). Eligibility criteria included age between 18 and 35 years of age, right-handedness, no history of neurological or psychological conditions, no limitations to upper limb mobility, and suitability for EEG procedures. Sample size for this experiment was not computed a-priori, but chosen to match and overtake that of other studies using similar analytical approaches (Häberle et al., 2023; Muhle-Karbe et al., 2021; Wolff et al., 2020).

Two participants were discarded because of poor EEG data quality (> 50% of trials dropped due to excessive noise) and technical issues during data collection. Exclusion criteria based on behavioral performance caused the removal of one participant due to accuracy below 60% in response to catch trials, and one participant because their mean delta time in the Joint task exceeded 3 standard deviations from the group mean, implying poor compliance to the experiment instructions (see below for the full explanation of the task and measures considered). Following exclusions, the final sample consisted of 36 participants (M_age_ = 24.56, ±4.86; 27 females and 9 males), all right-handed and with normal or corrected-to-normal vision acuity. Participants received a compensation of 30€ for their time.

## Materials

Each participant performed the task sitting comfortably at approximately 90 cm from a computer monitor (resolution 3840 x 2160, diagonal display length ∼69.47 cm, set on a 30Hz refresh rate). They were asked to use their right hand to draw on a graphic tablet oriented vertically (One by Wacom, model number CTL-472, with an active area of 15.2 x 9.5 cm). Stimuli presentation, collection of responses from both the keyboard and the tablet, as well as the interaction with the EEG recording software, were managed through the Psychopy toolbox, version 2022.1.3 (Peirce, 2007). The continuous trace produced by the pen on the graphic tablet was displayed live on the monitor, as a white dot (8 pixels radius) moving on a gray background, with a sampling rate of 30 Hz (i.e., the position of the dot was updated according to the new coordinates of the pen on the tablet every ∼33 ms). The entire surface of the graphic tablet was mapped on the right hemispace of the monitor, meaning that the left edge of the tablet corresponded to the vertical midline of the monitor. Participants started the experiment with extensive practice to familiarize with the experience of drawing on the tablet, and were encouraged to explore the limits of the drawing space to grow accustomed to the correspondence between the surface of the tablet and the monitor. A solid black line (4 pixels) was depicted along the midline, spanning 1200 pixels from the center of the monitor (11.81 degrees of visual angle). At the top and bottom of this line were two black circles of 100 pixels diameter (1 degree of visual angle), that we will refer to as starting and ending point, respectively (Figure 1).

**Figure 1.**
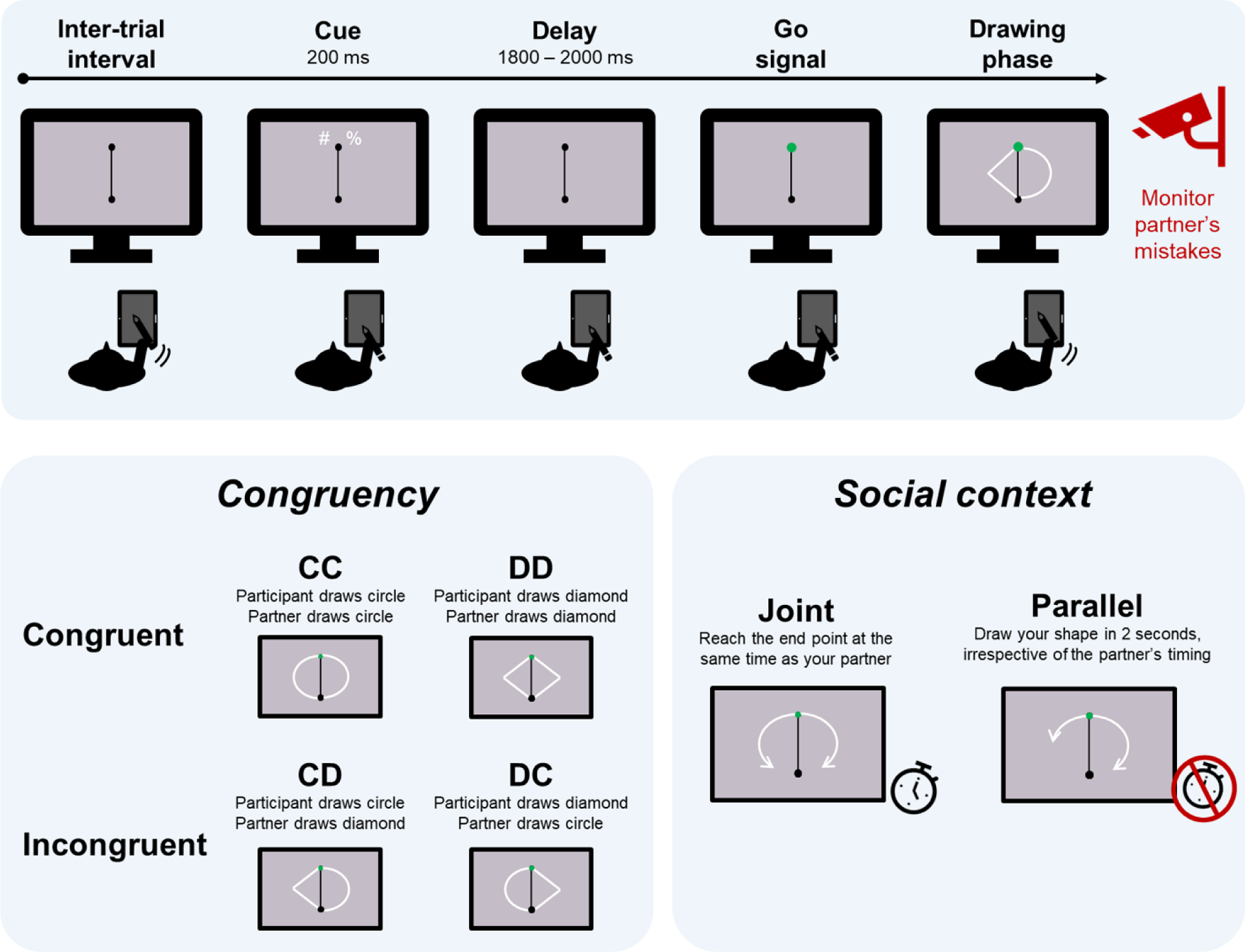
Behavioral Paradigm. *Note*. *Upper panel*: timeline of one example trial. The cue instructs the participant to draw a circle, while the partner will draw a diamond. The shapes are executed after the starting point turns green. *Bottom left panel*: Congruency manipulation. Depending on the combination of shapes drawn by the participant and the partner, trials could be congruent or incongruent. *Bottom right panel*: Social context manipulation. Participants carried out a Joint and a Parallel task, requiring to synchronize the drawing time with the partner or to draw consistently in 2 seconds, respectively.

Participants were informed that they would perform their task together with a partner. They would not meet or interact directly with the partner, but only see their drawings appear on the left side of the monitor, while their own drawing appeared on the right side. Drawings of both the participant and the virtual partner began at the starting point and reached the ending point. Both agents drew either a half circle or a half diamond, depending on the specific instructions for each trial. In reality, the drawings of the partner consisted of a set of pre-recorded hand-free drawn circles and half diamonds. This was done to maximize the feeling that these trajectories were the products of another human agent. However, their presentation was carefully controlled in order to match specific timing criteria (see below). To increase the perceived fluidity of the movement, the trajectories were displayed as thin white lines (5 pixels) originating from the starting point and progressively extending over time to reach the end point, with two new xy coordinates being added to the display on each screen refresh.

## Procedure

### Familiarization phase

All participants started the experiment with a familiarization phase to get used to drawing with the tablet, during which they were allowed to draw freely. When they felt comfortable with it, they moved on to the next phase, in which they learnt how to accurately draw the shapes required for the experiment. They were instructed to guide the tip of the pen in correspondence of the starting point, triggering the appearance of a cue above it, on the right-hand side. The cue indicated the shape to draw: these could be either half a circle or half a diamond, initiating at the starting point and reaching the end point at the bottom of the midline. The cue could be one of four symbols (#, $, %, &). Each participant was assigned one of 6 possible cues combinations. Namely, for the whole experiment two symbols indicated circles, and two symbols indicated diamonds. At the beginning of each block, one symbol for each shape was instructed to the participant, and the cue pairs kept alternating across blocks.

The cue stayed on display for 200 ms, followed by a cue-target interval with a duration uniformly distributed between 1800 to 2000 ms. Then, the starting point would change its color to green, prompting the participant to start drawing (i.e., go signal). During this first practice phase (i.e., *Shape familiarization*), a gray shaded area appeared on the monitor to guide the participant in their drawing, indicating the area boundaries within which the drawing would be considered accurate (i.e., an area of 100 pixels surrounding the perfect circle and diamond trajectories between the starting and the ending point). To make this learning phase even more effective, the leading tip of their drawn trajectory would be large (15 pixels) if its location was within the indicated shaded area, whereas it would be as small as the size of the remaining trailing trace if outside of the provided boundaries. Participants completed 6 of such familiarization trials, without any time constraints.

Once the expected shapes were learnt, participants completed 12 additional trials (6 for each shape) without the aid of the shaded area to guide them (i.e., *Execution practice*). They were instructed to replicate the same shapes they practiced in the previous step, and received analogous online feedback in the form of the pen tip diameter according to the same criteria. Here, they were additionally required to draw their shapes in approximately 2 seconds, and to initiate the movement as soon as possible when the starting point turned green. Feedback was provided after each trial, evaluating the drawing based on three criteria: 1) the trajectory was considered accurate if at least 60% of its coordinates fell within the ideal trajectory boundaries; 2) the total drawing duration was between 1.6 and 2.4 seconds; and 3) the starting time (i.e., interval between the starting point turning green and the tip of the pen leaving it) was below 800 ms.

Next, participants completed an *Observation practice* phase, during which they were not required to draw, but were introduced to the drawing performed by the virtual partner on the left side of the monitor. Analogously to the Execution practice, the trial started when the tip of the pen was placed within the starting point, this time triggering the appearance of a symbol above it on the left side. This indicated the movement that the virtual partner was about to perform. After the go signal and a delay spanning from 400 to 600 ms, one of the pre-drawn trajectories started being displayed, for a total duration ranging from 1.6 and 2.4 seconds. In 4 trials out of 12, the trajectory being displayed did not match the identity of the presented cue (i.e., *catch trials*). Participants were instructed to monitor the performance of their virtual partner, and to press the ‘b’ key on the keyboard as soon as they noticed a mistake in the shape being drawn. The goal of introducing catch trials was to ensure that attention was allocated to the action of the partner, and that its cued identity was retained during the whole delay interval.

Once these task subcomponents had been practiced individually, they were combined and performed together. Namely, for each trial, two symbols would appear as cues, one indicating a shape for the participant and one for the partner. These could be the same (i.e., congruent trials), or two opposite shapes in the case of incongruent trials. After the go signal, the goal of the participant was to draw their shape, and also to monitor the drawing of the partner to detect potential catch trials.

### Social context manipulation: practices and main tasks

The core social manipulation in our experiment consisted of a Joint and a Parallel conditions, administered in a block design to all participants, with their order counterbalanced. The two social contexts were designed to be as similar as possible in terms of visual input, amount and relevance of the information provided to the participant in each trial, motor requirements, and overall movement durations. What distinguished the two conditions was the extent to which the participant had to online adjust their movement to that of the partner. Namely, in the Joint social context, the requirement for participants was to draw accurately, and to synchronize their drawing time to that of the partner, with the goal of reaching the end point simultaneously. On the contrary, in the Parallel context participants had to draw accurate trajectories, trying to keep their timing consistent to approximately 2 seconds, irrespective of the drawing time of the partner. In other words, the specific kinematic properties of the partner’s movement were behaviorally relevant for the participants only during the Joint task, as they had to adjust their behavior to the speed of the partner’s movement, whereas in the Parallel task only the identity of the shape drawn by the partner had to be monitored to detect catch trials (Figure 1).

Each of the two conditions was preceded by a task-specific practice phase. Blocks of 12 trials (3 for each combination of shapes between participant and partner, namely both circles, both diamonds, circle and diamond, diamond and circle) were administered. Two trials in each block were catch trials (i.e., the partner drew the opposite shape to that indicated by their cue). The performance in these practice trials was again measured according to 3 criteria: 1) the trajectory was considered accurate if at least 60% of its coordinates fell within the ideal trajectory boundaries; 2) the starting time of the participant’s drawing was within 800 ms from the go signal; if the starting time exceed 800 ms, the trial was interrupted and considered an error; 3) the drawing time was in accordance to the task goals. More specifically, in the Joint context, a trial was considered good with respect to timing if the difference (i.e., delta) between the moment the participant and the partner reached the end point was below 200 ms. Small deltas indicate good synchronization to the movement speed of the partner. On the contrary, the Parallel context required participants to be consistent in their own timing, and trials were considered timely if the drawing was within 200 ms from the instructed 2 seconds duration. After each sequence of 12 practice trials, the performance in the block was evaluated. If across trials the average trajectory accuracy was above 60%, there were no more than 2 trials with slow starting time, at least one catch trial was correctly detected, and the task-specific timing requirement was fulfilled, the practice ended and the main task would start. Otherwise, another practice block would start, up to a maximum of 8 blocks. On average, 2.6 (±1.85) blocks were completed for the Joint context, and 2.4 (±2.02) for the Parallel context. The number of practice blocks needed for the two tasks did not differ significantly across participants (W = 182.5, *p* = 0.65, CLES = 0.57).

Following the successful completion of the task-specific practice, 6 blocks of 48 trials were completed for each of the two social contexts. Each participant completed 60 trials for each context and shape pairs combination, 12 of which were catch trials, resulting in a total of 576 trials. The trial structure was identical to the practice, with the only difference that feedback was provided only for missed catch trials (as a message stating that ‘your partner drew the wrong shape!’) and for trials with slow starting time (‘You waited too long to start drawing. Try to start drawing as soon as the starting point turns green!’). Trials with a slow start were immediately interrupted and repeated at the end of the block (maximum once). In each block of the main task, 6 were catch trials, to ensure in both social contexts participants retained the cue identity for the partner’s shape throughout the whole trial, and allocated comparable attention to the drawing of the partner. Feedback was provided at the end of each block as the average accuracy in terms of trajectory, timing, detection of partner’s mistakes.

Crucially, timing parameters were adjusted carefully during the Joint and Parallel contexts, in order to characterize the different task requirements. In the Joint task, the movement duration of the partner was uniformly distributed from 1.8 to 2.2 seconds. In the Parallel task, the movement duration of the partner was sampled from a slightly larger distribution, spanning from 1.6 to 2.4 seconds. The reason for this discrepancy was to ensure participants could not rely on the movement duration of the partner to fulfill the timing requirements of the Parallel task (i.e., in some trials the partner would be too fast or too slow with respect to the accepted timing criterium), and therefore ensuring a difference in how the two tasks were approached and performed, while keeping all other factors equal.

Throughout the whole experiment, another timing parameter was updated after each block. The participant’s starting times (i.e., time between the go signal and the pen tip leaving the starting point) were averaged at the end of each block. This value was then used in the subsequent block to adjust the starting time of the partner’s movement, sampled from a uniform distribution centered on the averaged starting time of the participant, and spanning ±100 ms. While keeping constant for the whole experiment the maximum allowed starting time of 800 ms, this adaptive starting time for the partner’s movement 1) guaranteed that the partner’s performance was credibly attributed to another human agent, and 2) reduced the likelihood of participants waiting for the partner’s drawing to start, in order to detect potential catch trials, and then proceed to execute their own movement. This setup ensured that drawing and monitoring of the partner’s performance happened simultaneously.

After the preparation and the EEG cap set-up (∼40 minutes), the total duration of the experiment, including the familiarization phase, the two task-specific practices, and the two main tasks, was of approximately 90 minutes. Participants could take a self-paced break between each block of the main task, and a longer break was encouraged after the end of the first main task, in order to allow participants to relax before learning and practicing the second task. At the end of the experimental phase, the EEG cap was removed and a short debriefing was provided to the participants.

## Experimental design and behavioral analyses

The experiment consisted of the within-subject factor Social context (Joint vs Parallel) and the within factor Combination, with four level corresponding to the crossing of the shape drawn by the participant and the shape draw by the partner (CC: both agents draw circles, DD: both agents draw diamonds, CD: the participant draws a circle and the partner a diamond, DC: the participant draws a diamond and the partner a circle; the first letter always indicates the shape of the participant, the second the shape of the partner). Crucially, the four combinations could be grouped for subsequent analyses in the factor Congruency, with the two levels Congruent (combinations CC and DD) and Incongruent (combinations CD and DC).

With respect to general behavioral performance in the two social contexts, we compared some indicators to evaluate whether participants approached the two tasks differently. First, we compared accuracy in response to catch trials between the two contexts by means of a non-parametric Wilcoxon signed-rank test (due to violated normality assumption, Shapiro-Wilk test *p* < 0.05). Then, we quantified whether the timing requirements of the two contexts were correctly implemented. For the Joint task, we expected average delta times to not differ significantly from 0, hence we tested for this comparison with a one-sample t-test. Analogously, in the Parallel task participants were instructed to draw consistently in 2 seconds, thus we compared the average duration of their movement (from the moment the pen left the starting point, to the movement it reached it) with this instructed duration. All statistical comparisons were carried out with the Python toolbox Pingouin version 0.5.3 (Vallat, 2018) unless otherwise specified.

## Trajectory analyses

With respect to overt motor behavior, our main goal was to quantify the visuomotor interference elicited by the partner’s movement on the drawing of the participant, and to investigate whether this was influenced by the social context manipulation. Trajectories were recorded as the collection of xy coordinates covered by the tip of the participant’s pen from the moment it left the starting point (i.e., beginning of the drawing action) to the moment it reached the ending point. Only trajectories recorded during non-catch trials, extending fully from the starting to the ending point were used for this analysis.

As one new pen coordinate was sampled at the refresh rate (30 Hz, one sample each ∼33 ms), trajectories recorded in each trial had different number of data points, depending on the overall drawing duration. Therefore, the first step to make trajectories comparable across trials and conditions was to interpolate and resample them to the same number of data points. For each trajectory (i.e., each trial) we fitted a cubic spline (with the default settings of the function interpolate.CubicSpline() from the scipy package), and sampled 100 points from it. Next, we visually inspected all the interpolated trajectories and discarded those in which the participant made gross errors (e.g., the pen slipped from their hand resulting in a meaningless scribble). On average, gross errors were identified in 5.45 (±6.88) trials per participant. Notably, we confirmed that this trimming procedure did not result in discarding more trials in any specific experimental condition, by submitting the count of discarded trials for each participant and condition to a Generalized Linear model with factors Social context and Congruency, using a Poisson distribution and Sum contrast coding (The jamovi project, 2023). We found no significant effect (all ps > 0.21), indicating that participants did not commit more gross errors in any specific experimental condition.

The visual inspection implemented was agnostic with respect to the shape indicated by the cue. In other words, we did not visually discard trajectories based on whether the drawn shape matched the instructed one or not. To objectively quantify such swap errors, for each trajectory we identified the rightmost coordinate point, and used it as knot to fit a linear and a quadratic univariate spline (interpolate.LSQUnivariateSpline from the scipy package, with an order or 1 and 2, respectively). Practically, we fitted two straight lines or two curves between the identified rightmost point, and the first and last point of the trajectory. Linear fits should approximate diamonds better, whereas quadratic fits were expected to better capture circles. Then, for each trial we checked which of the two fits performed better (i.e., had lower residuals) and marked as swap errors trajectories in which the fit inconsistent with the instructed cue had lower residual values. This resulted in an average of 4.22 (±7.43) trials marked as swap errors per participant. Again, we tested whether participants committed more swap errors in any experimental condition by means of a Poisson Generalized Linear Model, with factors Social context and Congruency (The jamovi project, 2023). We found no significant effect (all ps > 0.150). Since it cannot be disentangled whether swap errors were due to an inaccurate retention of the cue identity, or were the result of visuomotor interference, and since they did not differ in number across conditions, we discarded these trials from subsequent analyses. Overall, we retained for analyses an average of 462.08 (±15.11) trajectories per participant (115.52 ±4.24 for each Social context x Congruency condition).

Visuomotor interference in continuous movements has been defined in previous work as the variability in movement trajectories (Kilner et al., 2003). To quantify visuomotor interference in our task, we extracted for each trial an index of distortion of the drawn trajectory from a condition-nonspecific template. For each participant, we first computed a template circle and a template diamond, by averaging all instances of each of the two shapes, across all experimental conditions. Then, we measured the area subtended by the trajectory drawn in each individual trial and the corresponding template shape. This resulted in a single value per trial, indicating how distant the trajectory was from the participant-specific average shape template, with larger values indicating larger deviations from the template (Figure 4, left panel). Because the resulting distribution of these values severely violated normality, we applied a Box-Cox transformation (Atkinson et al., 2021; Pek et al., 2018), as implemented in the scipy.stats package (Virtanen et al., 2020). The optimal lambda was estimated to be 0.336, and the transformed data successfully approximated normality (skewness = -0.037, kurtosis = -0.164). Therefore, we fitted this Box-Cox transformed distribution to a Linear Mixed model using the lme4 package in R (Bates et al., 2014). We included as fixed effects Social context, Congruency, and their interaction; and we included a random intercept for each participant (in lme4 notation: *Area ∼ SocialContext * Congruency + (1 | Subject)*), setting a Sum contrast coding for both predictors (-1, 1). Notably, adding random slopes to the model equation resulted in failure of model convergence, therefore we implemented the simple intercept model. Visual inspection of the residuals did not reveal a deviation from normality. P-values were computed with the Satterthwaite-corrected degrees of freedom (Kuznetsova et al., 2017).

Additionally, based on the Dyadic Motor Plan framework and previous evidence linked to it (Clarke et al., 2019; Sacheli et al., 2018, 2019), we formulated also the more specific and directional hypothesis of larger distortions in incongruent trials of the Parallel task. To test it, we computed the Congruency Effect (i.e., mean distortion in incongruent trials – mean distortion in congruent trials), separately for each of the two tasks. We then compared the Congruency Effects across Tasks with a paired-samples one-tailed t-test, assuming larger values for the Parallel task.

To ensure distortions in the drawn trajectories are not caused by confounding factors other than the target experimental manipulations, we performed two additional control analyses. First, we added to the model described in the previous paragraph an additional fixed effect, namely the centered drawing time (in lme4 notation: *Area ∼ SocialContext * Congruency * DrawingTime + (1 | Subject)*). The rationale for this control analysis was to rule out that larger distortions could be due to faster drawing performance, akin to a speed-accuracy tradeoff. Second, we tested for the effect of the order in which the two tasks were performed. Namely, participants performed the two tasks in a blocked fashion, with the order of the two tasks counterbalanced across participants. With this control analysis we aimed at checking that the visuomotor interference was not systematically influenced by the order in which the two tasks were administered. To this goal, we fitted the model *Area ∼ SocialContext * Congruency * TaskOrder + (1 | Subject)*.

## EEG recordings and preprocessing

Electrophysiological data were recorded with a Biosemi ActiveTwo system, with 64 Ag-AgCl electrodes arranged in accordance to the international 10-20 system (Klem et al., 1999). The setup included a Common Mode Sense - Driven Right Leg (CMS-DRL) electrode pair, two external electrodes attached to the left and right mastoids, and four electrodes (two to the outer canthi of both eyes, one above and one below the left eye) used to monitor horizontal and vertical eye movements. Data were recorded at a sampling rate of 1024 Hz, and we aimed at keeping impedance below 10 kΩ.

All preprocessing steps were carried out using the MNE Python toolbox version 1.3.1 (Gramfort, 2013). First, to achieve the maximum signal-to-noise ratio provided by the CMS-DRL setup, we initially re-reference the data to the electrode POz (spatially located in the cap between CMS and DRL). Then, we implemented a bandpass FIR filter on the continuous data, in the frequency interval 0.1 – 40 Hz (Hamming window with 0.0194 passband ripple and 53 dB stopband attenuation, with lower and upper transition bandwidth of 0.1 and 10 Hz, respectively). Next, we used the NoisyChannels function, available in the toolbox PyPrep version 0.4.2 (Appelhoff et al., 2022; Bigdely-Shamlo et al., 2015) to detect bad channels in the continuous recordings. We adopted the default parameters of the toolbox and searched for bad channels according to the whole range of methods available, namely: low signal-to-noise ratio, channels with poor correlation with the surrounding ones, channels containing abnormally high amplitudes, excessive high frequency noise, flat or near-flat values, and channels poorly predicted by the others (*ransac* approach). This comprehensive procedure resulted in 3.64 (±1.83) channels being flagged as bad. Note that for all participants the channel POz was indicated as flat due to it being used as reference.

Our main interest was to test for the preparatory neural representations associated to dyadic movements. Therefore, we focused our analyses in the delay period between the cue and the go signal. The continuous data was epoched time-locked to the onset of the cue (initially from -500 to 3000 ms), linearly detrended, and downsampled to 128 Hz. To remove electrophysiological activity associated with eye movements, an Independent Component Analysis (ICA) was performed and components were selected to be discarded when they correlated with the activity recorded by the electrooculogram located above the left eye (MNE function ica.find_bads_eog()). On average, 1.25 (±0.6) components were discarded. After re-referencing to the average of all electrodes, we proceeded to discard trials in which the 150 µV threshold was exceeded (on average, 17.08 ±25.45 trials per participant). Finally, epochs were baseline corrected to the average signal in the time window 200-0 ms preceding the cue onset.

As the focus of our hypotheses was the delay period following the cue and preceding the go signal, both regular and catch trials were included in the analyses. Catch trials that were not correctly identified, and regular trials that were indicated to be catch trials were excluded from the analyses. For each individual condition (crossing of the factors Social context and Congruency), we retained an average of 139.74 (±6.36) trials.

## EEG decoding analyses

### Decoding action combinations in Congruent and Incongruent trials

In line with the Dyadic Motor Plan hypothesis (Sacheli et al., 2018), we expected that, during the preparation interval of the Joint condition, the upcoming participant’s and partner’s movements would become integrated in a conjoined representation. We reasoned that this would result in low decoding accuracies when trying to discriminate incongruent combinations (CD vs DC), as both would include the same set of two movements. On the contrary, information concerning the two movements are thought to be maintained during Parallel actions in a less intertwined fashion, leading to higher discriminability. Given the complete overlap between the two upcoming actions in the Congruent combination pairs, we expected high discriminability in both Social contexts. Therefore, to test this hypothesis, we performed 2-way classifications on the broadband EEG data, with the goal of classifying the two possible movements combinations (i.e., CD vs DC and CC vs DD), separately for the two contexts. The classification procedure was carried out using the Scikit-learn Python toolbox version 1.0.2 (Pedregosa et al., 2011) and MNE Python version 1.3.1 (Gramfort, 2013).

To increase signal-to-noise ratio in the input for the classifier, we created *supertrials* by averaging together randomly sampled sets of four trials belonging to the same category (Grootswagers et al., 2017). To minimize the influence of the visual features of the cue on the classifier, for each supertrial we averaged together trials belonging to the same experimental condition, but randomly sampled in equal number from each of the two cue pairs (e.g., one supertrial for the condition CC would be the result of averaging together two CC trials with cue pair 1, and two CC trials with cue pair 2). For each classification, we created 10 supertrials for each condition. The mean activity was then subtracted from each supertrial and each channel separately, to normalize the voltage fluctuation over time (Muhle-Karbe et al., 2021; Wolff et al., 2020).

Instead of performing decoding separately for each time point, we implemented spatiotemporal decoding, capturing multivariate information encoded not only in spatial patterns, but also in their temporal evolution (Muhle-Karbe et al., 2021; Wolff et al., 2020). With this approach, both spatial and temporal features of the neural activity are used for classification within participant, with the advantages of improving decoding accuracy and being insensitive to potential differences in decoding latencies across participants. We were interested in the neural representations arising during the delay period, thus anticipating the overt movement. Therefore, our time window of interest covered the interval between the onset of the cue and the subsequent 2000 ms (i.e., the shortest possible delay interval, accounting for its variability). We averaged the data from these 2-seconds window in 10 bins of 200 ms. As a result, each single supertrial was treated as a discrete event, with 640 features (i.e., 10 binned-values for each of the 64 electrodes).

These binned supertrials were used as input to train and test linear discriminant analysis (LDA) classifiers, with a least square solver and automatic shrinkage based on the Ledoit-Wolf lemma. We used a Stratified 5-Fold cross-validation approach. This means that the pool of twenty supertrials entering each classification was split 5 times in non-overlapping training sets of 16 supertrials, and testing sets of 4 supertrials, equally representing the two classes, and the classification accuracy was averaged across the 5 folds.

Importantly, to avoid biases due to the random sampling of trials in creating supertrials, for each contrast of interest we repeated the whole classification pipeline for 100 permutations (i.e., we generated supertrials and performed the 5-Fold cross validated classification for 100 times) and we averaged the results (Figure 2).

**Figure 2.**
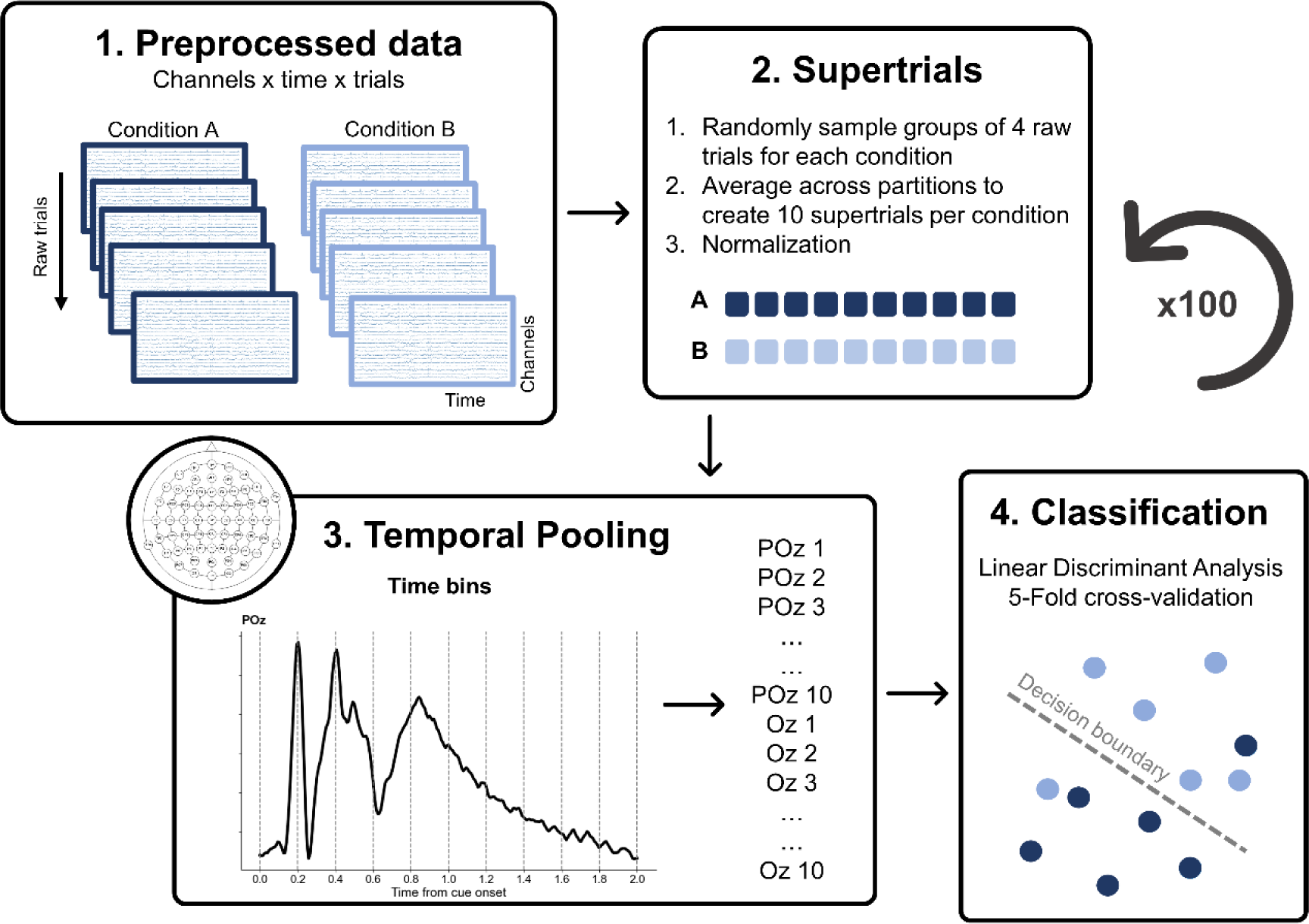
Spatiotemporal Decoding. *Note.* Epoched data after preprocessing (1) are averaged together in random partitions of the same conditions to create supertrials (2). Time courses for each of the 64 channels are pooled in time, averaging voltages in ten time-bins (3). The resulting 640 values are used as features to train and test an LDA classifier (4). The whole procedure is repeated 100 times.

### Decoding own and partner’s movements

Additionally, we tested whether the two instructed movements were decodable separately during the delay period. Therefore, we adopted the same spatiotemporal approach to classify the upcoming movement of the participant, and the upcoming partner’s movement, across Congruency levels and separately for the two Social contexts. In other words, we aimed at classifying whether the participant was instructed to draw a circle or a diamond (conditions CC and CD vs conditions DC and DD), and, separately, if the partner’s instruction was to draw a circle or a diamond (conditions CC and DC vs conditions CD and DD).

The only difference from the classification pipeline explained above was in the creation of supertrials. To classify the upcoming movement of the participant, we created supertrials averaging together one individual trial from each of the two experimental conditions in which the participant had to draw a specific shape, and one for each physical cue identity. For example, to create one supertrial for the condition “participant was instructed to draw a circle”, we averaged together one trial from condition CC (cue pair 1), one trial from condition CC (cue pair 2), one trial from condition CD (cue pair 1), and one trial from condition CD (cue pair 2). This resulted in the same total number of trials (i.e., four) contributing to the formation of one single supertrial, as in the previous contrasts.

Analogous 5-Fold cross-validated LDA classifications were performed, repeating the random sampling for the supertrials creation for 100 permutations.

### Statistical testing of spatiotemporal decoding accuracies

The spatiotemporal classification approach described resulted in one value per participant for each contrast. We evaluated whether the resulting decoding accuracies were significantly higher than chance level (50% for 2-way classifications) via one-tailed one-sample t-tests.

Our main hypothesis was that decoding of incongruent movements combinations would be lower during Joint compared to Parallel task. To directly test for this, we performed a two-tailed paired-samples t-test contrasting decoding accuracies for Incongruent conditions (CD vs DC) between Joint and Parallel task. For completeness, we also compared accuracies in decoding congruent action combinations across social contexts. Finally, with respect to the decoding of own and partner’s movements, we compared whether decoding accuracies were significantly above chance via one-tailed one-sample t-tests, and were different between the two contexts, by means of two-tailed paired-samples t-tests.

### Spatial decoding

To have an understanding of which channels contributed most to the decoding obtained with our spatiotemporal approach, we repeated the same set of two-way classifications in a spatial fashion. Instead of treating space (i.e., channels) and time as features for the LDA classifiers, we performed the decoding on the time-resolved voltage values, separately for each channel. This analysis results in one decoding accuracy for each channel, participant, and contrast of interest, therefore allowing to display topographies of classification accuracies. For exploratory purposes, we conducted statistical testing to detect clusters of electrodes with decoding accuracies significantly larger than chance. We used a cluster-based permutation approach (Maris & Oostenveld, 2007; Sassenhagen & Draschkow, 2019), clustering across neighboring channels based on Delauney triangulation.

## Results

### General behavioral measures

We computed some indicators of behavioral performance to investigate compliance to the experiment instructions. First, we compared accuracy in catch trials detection between Joint and Parallel tasks. We designed the experiment with the goal of equating the attentional deployment to the partner’s drawing across the two conditions. Participants were highly accurate in identifying errors in partner’s trajectories during both Joint (Mean = 0.94, sd = 0.05) and Parallel task (Mean = 0.92, sd = 0.06), although slightly better during the Joint task (W = 162.0, *p* = 0.035, CLES = 0.62).

Next, we aimed at testing whether the timing requirements imposed by the task were attended correctly by participants. In the Joint task, we expected participants to align their timing with the movement duration of the partner’s, resulting in very small intervals between the moment theirs and the partner’s trajectories reached the ending point (i.e., delta times). As expected, we found that on average the delta time (Mean = 0.01, sd = 0.051) was not significantly different from 0 (t_35_ = 1.08, p = 0.28, d = 0.18, BF_10_ = 0.31). The total duration of the movement time was not analyzed statistically for the Joint context (Mean = 2.07, sd = 0.06), as it was biased by the discrepancy between the participant’s starting time and the (adaptive) starting time of the partner.

With respect to drawing time in the Parallel task, participants were instructed to complete their trajectory in 2 seconds. However, they showed a tendency to systematically shorten their drawing time (t_35_ = -2.71, p = 0.01, d = 0.18), resulting in an average drawing time of 1.93 seconds (±0.15).

Overall, these results support the effectiveness of our social manipulation, indicating that participants understood the timing requirements of the two tasks and behaved accordingly, although they were more successful in complying with the requirements of the Joint task.

### Visuomotor interference is larger in the Parallel context

Our main hypothesis concerning visuomotor interference in the drawn trajectories was that an incongruent movement should result in larger distortions during the Parallel task. We predicted that, in the Joint task, the active anticipation of the partner’s drawing, and its integration into a Dyadic Motor Plan, would drastically reduce the distortion in the participant’s drawing, in line with other experimental findings (Rocca et al., 2023; Rocca & Cavallo, 2020; Sacheli et al., 2018).

The results of the fitted Linear Mixed Model revealed a significant effect of Social context (β = -0.53, 95% CI = [-1.02, -0.05], t_16629_ = -2.17, *p* = 0.030), indicating that trajectories were deviating more from the template in the Parallel compared to Joint task. The effect of Congruency approached but did not cross the significance threshold (β = -0.45, 95% CI = [- 0.94, 0.03], t_16629_ = -1.84, *p* = 0.065), hinting at larger distortions in incongruent trials. Finally, the interaction effect also approached but did not reach the significance threshold (β = 0.45, 95% CI = [-0.003, 0.93], t_16629_ = 1.82, *p* = 0.069).

To have a more direct test of our effect of interest, we computed the Congruency Effect for each of the two contexts (see methods). The one-tailed t-test revealed significantly larger distortions in the Parallel compared to Joint task (t_35_ = -1.88, *p* = 0.034, d = 0.43, BF_10_ = 1.73). Despite the small effect size, these results are in line with the hypothesis that integrating the partner’s action into a Dyadic Motor Plan reduces the interfering effect of observing an incongruent movement (Figure 4).

**Figure 3.**
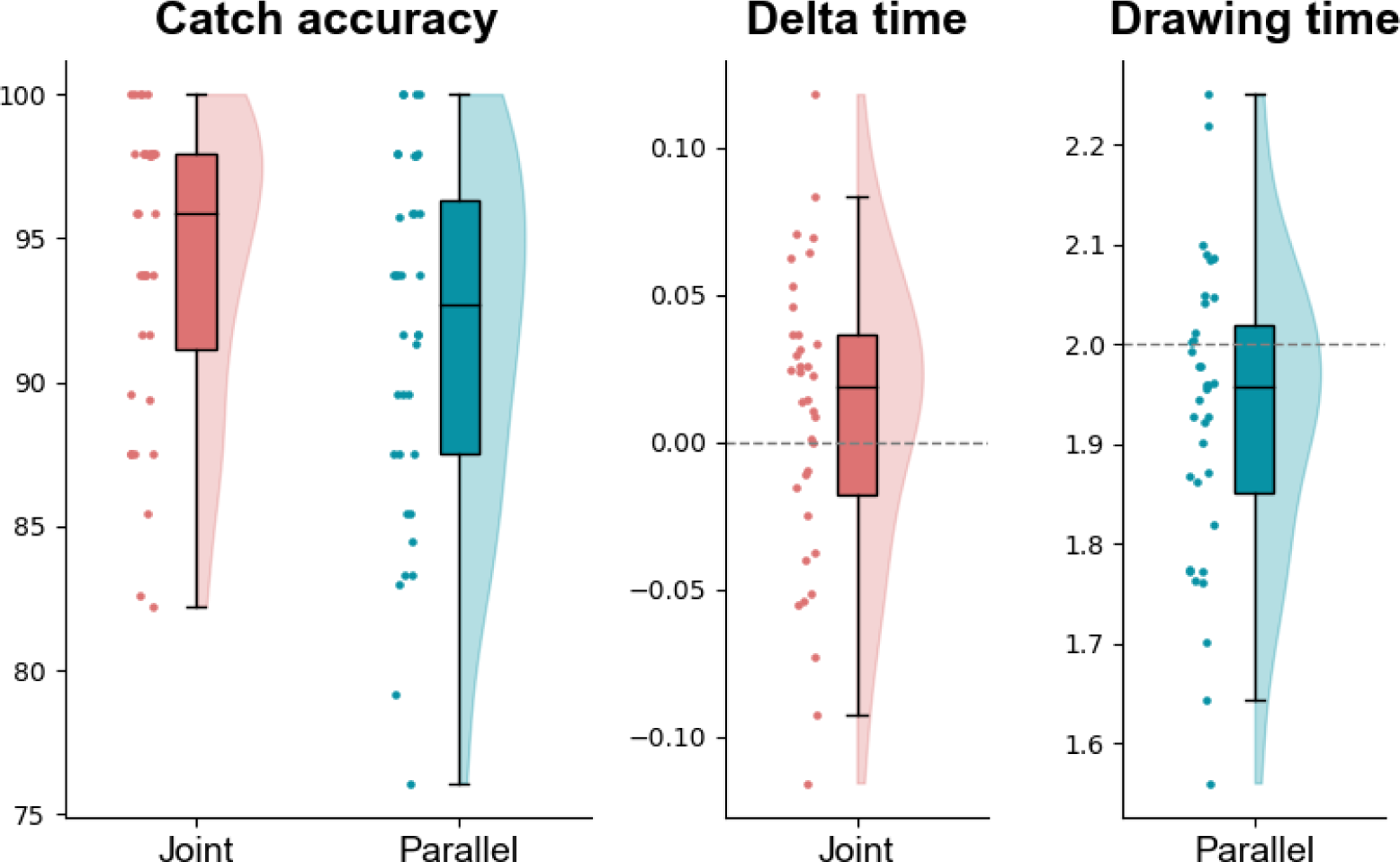
Indicators of Behavioral Performance. *Note.* In each boxplot, the black line inside the box indicates the second quartile (median) of the distribution (n = 36). The bounds of the boxes depict the first and third quartiles of the distribution. Whiskers denote the 1.5 interquartile range of the lower and upper quartile. On the left side of each boxplot, dots represent individual scores, whereas on the right side their density is displayed as a half violin plot.

**Figure 4.**
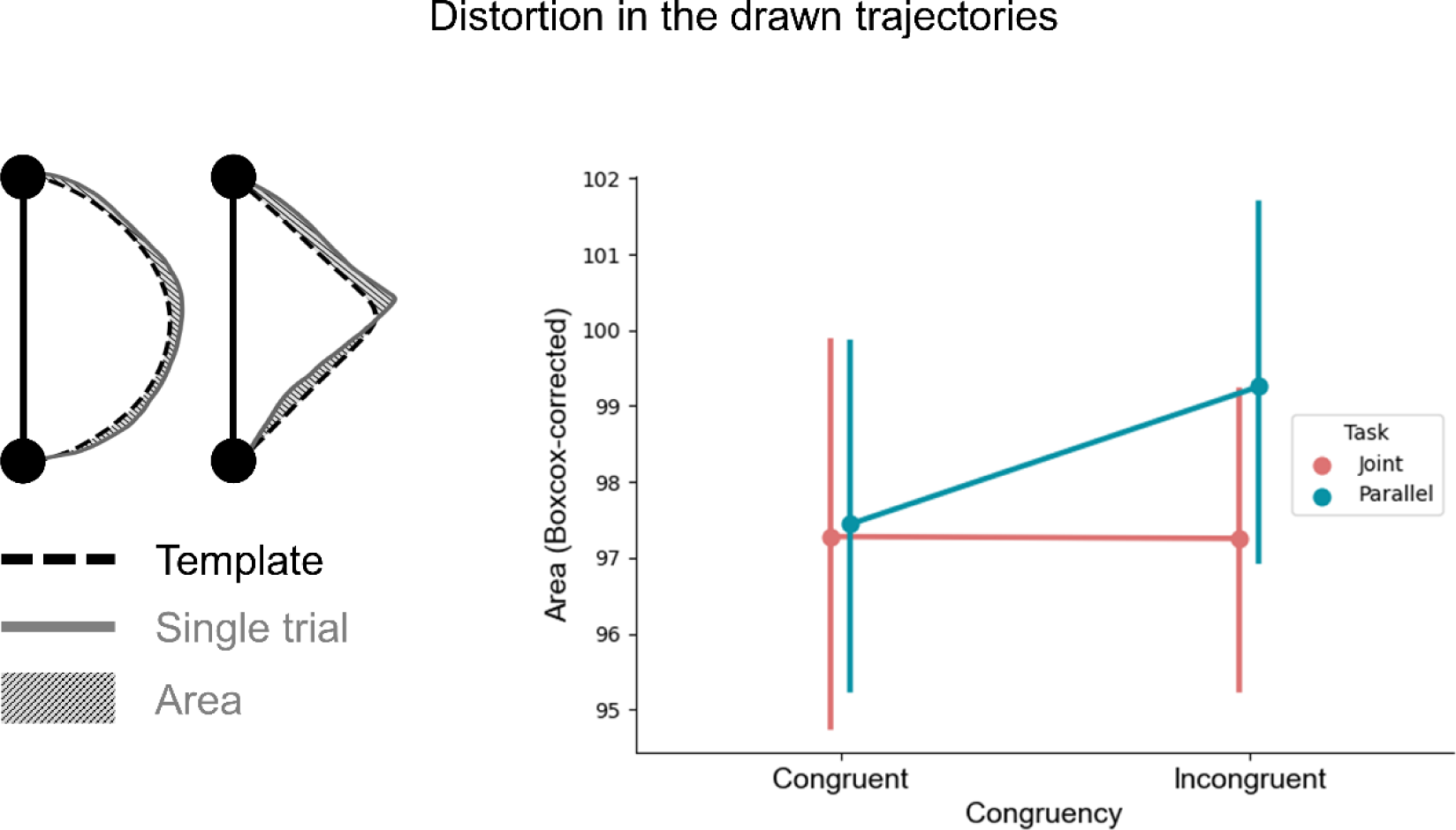
Visuomotor Interference Results. *Note*. *Left panel*: VMI is quantified as the area between the average shape templates for the participant, and each of the drawn trajectories. *Right panel*: Boxcox corrected Area values, for each Social context * Congruency condition. Error bars depict the confidence interval.

We fitted two additional models as control analyses. First, we extended our model to include the centered factor Drawing Time and its interactions with the factors Social context and Congruency. The rationale of fitting this model was accounting for a potential effect of the speed of drawing on the distortion in the trajectories. The results of this model were analogous to the previous analysis with respect to the main effects of Social context, Congruency and their interaction. Additionally, we found a significant interaction of Congruency and Drawing Time (β = -0.78, 95% CI =-1.27, -0.30], t_16625_ = -3.18, *p* = 0.001). Notably, this interaction indicated that slower movements were associated with larger distortions in incongruent trials, discarding the possibility of a speed-accuracy trade-off between movement duration and variability in drawn trajectories (Supplementary Materials Figure S1). Next, we extended our original model to include the factor Task Order and its interaction with Social context and Congruency. While this model replicated again the main findings concerning the experimental manipulations of interest, the effect of Task Order and all the related interactions were not significant. We took these results as evidence that the interfering effect of incongruent movements during the Parallel task were not associated with the order in which the participants performed the tasks, nor could be attributed to an increased variability due to faster drawing times.

### Social context influences the neural representation of upcoming movements

To test whether the Joint social context induced the formation of a Dyadic Motor Plan, in which the movements of the two agents are tightly integrated, we aimed at classifying conditions pairs, separately for Joint and Parallel task (Figure 5). First, we tested whether the classifier could pick up differences between the two incongruent conditions (i.e., CD vs DC). We reasoned that, if the two actions are represented intertwined in the Joint condition as proposed, it should be more difficult for the classifier to tease these two conditions apart. We found high above chance decoding accuracies in the Parallel task (t_35_ = 5.06, *p* < 0.01, d = 0.84, BF_10_ > 100), and smaller but still significant decoding accuracies in the Joint task (t_35_ = 1.78, *p* = 0.04, d = 0.30, BF_10_ = 1.50). Crucially, we observed a significant difference between Joint and Parallel tasks in the decoding accuracies of incongruent combinations (t_35_ = 2.74, *p* < 0.01, d = 0.59, BF_10_ = 4.37).

**Figure 5.**
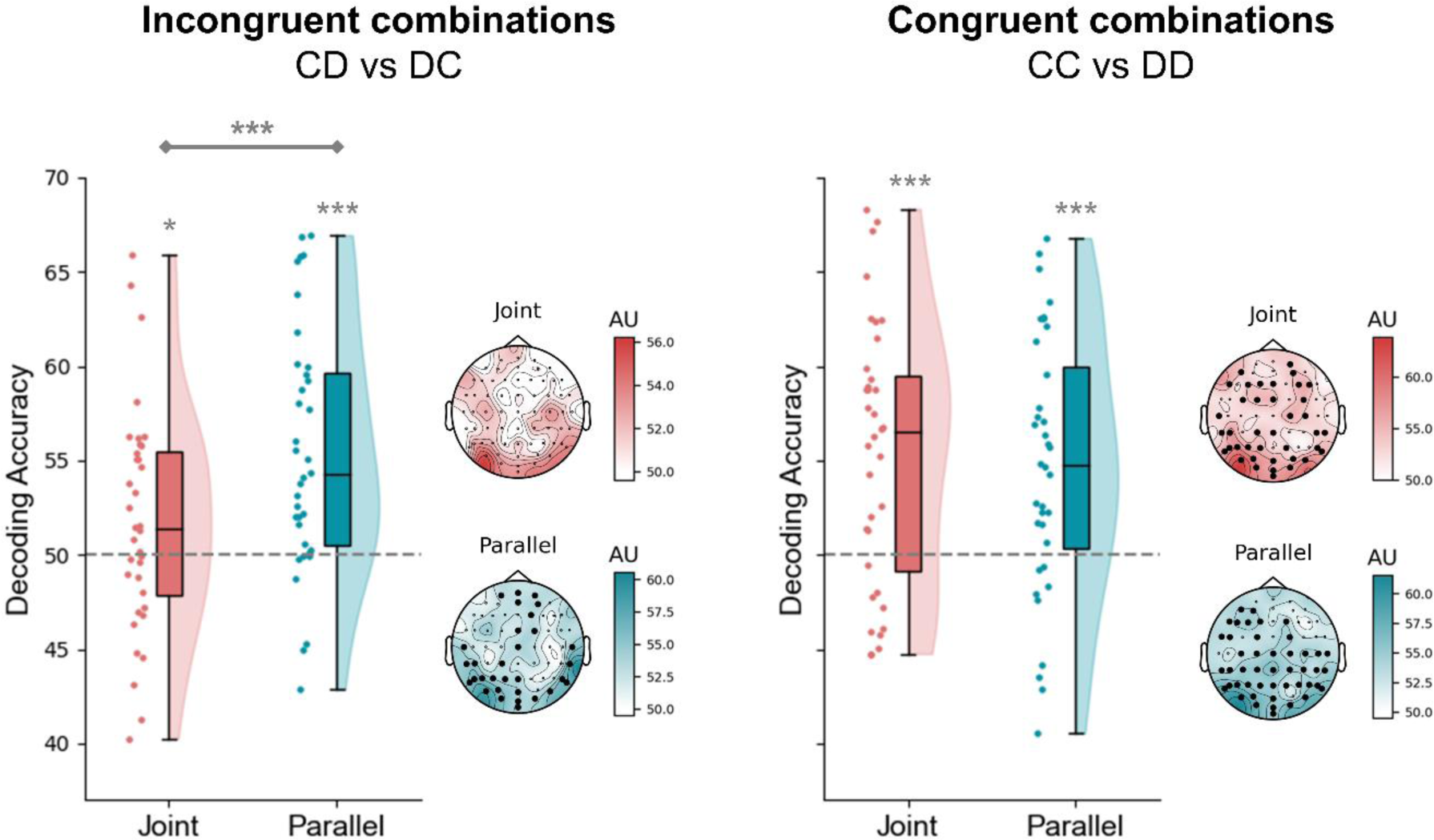
Decoding of Incongruent and Congruent Combinations. *Note. Left panel*: accuracies resulting from the decoding of the conditions CD vs DC in Joint and Parallel task. The dashed horizontal grey line indicates the chance level (50%). In each boxplot, the black line inside the box indicates the second quartile (median) of the distribution (n = 36). The bounds of the boxes depict the first and third quartiles of the distribution. Whiskers denote the 1.5 interquartile range of the lower and upper quartile. On the left side of each boxplot, dots represent individual scores, whereas on the right side their density is displayed as a half violin plot. For visualization purposes, the topographies report the decoding accuracies for each of the 64 channels, separately for each task. Thicker dots indicate channels belonging to spatial clusters with significantly above chance decoding accuracy. *Right panel*: accuracies resulting from the decoding of the conditions CC vs DD in Joint and Parallel task. Plotting conventions are the same as for the left panel. *: *p* < 0.05, ***: *p* < 0.001

With respect to discriminating between congruent movement combinations (i.e., CC vs DD), the two movement combinations were successfully classified above chance level in both the Joint (t_35_ = 4.64, *p* < 0.01, d = 0.77, BF_10_ > 100) and Parallel task (t_35_ = 4.16, *p* < 0.01, d = 0.69, BF_10_ > 100). The observed decoding accuracies did not differ between the two social contexts (t_35_ = 0.43, *p* = 0.67, d = 0.09, BF_10_ = 0.19).

For qualitative exploration, we implemented the same decoding contrasts also at the spatial level, to identify those electrodes carrying most information. With the exception of the incongruent combinations in Joint task, the other contrasts yielded electrodes clusters of above-chance accuracy covering mostly occipito-parietal and frontal regions.

One property of the classification approach we adopted is that it is agnostic to what aspect of the data is informing the classifying algorithm. Namely, in our comparisons the classifier might be picking up information encoding the upcoming participant’s movement, as well as the partner’s movement. Therefore, to better disentangle the individual contributions of these two sources of information, we carried out two additional classification analyses, aimed at discriminating the upcoming participant’s movement (Circle: CC and CD vs Diamond: DC and DD), and the upcoming partner’s movement (Circle: CC and DC vs Diamond: CD and DD), across Congruency levels (Figure 6).

**Figure 6.**
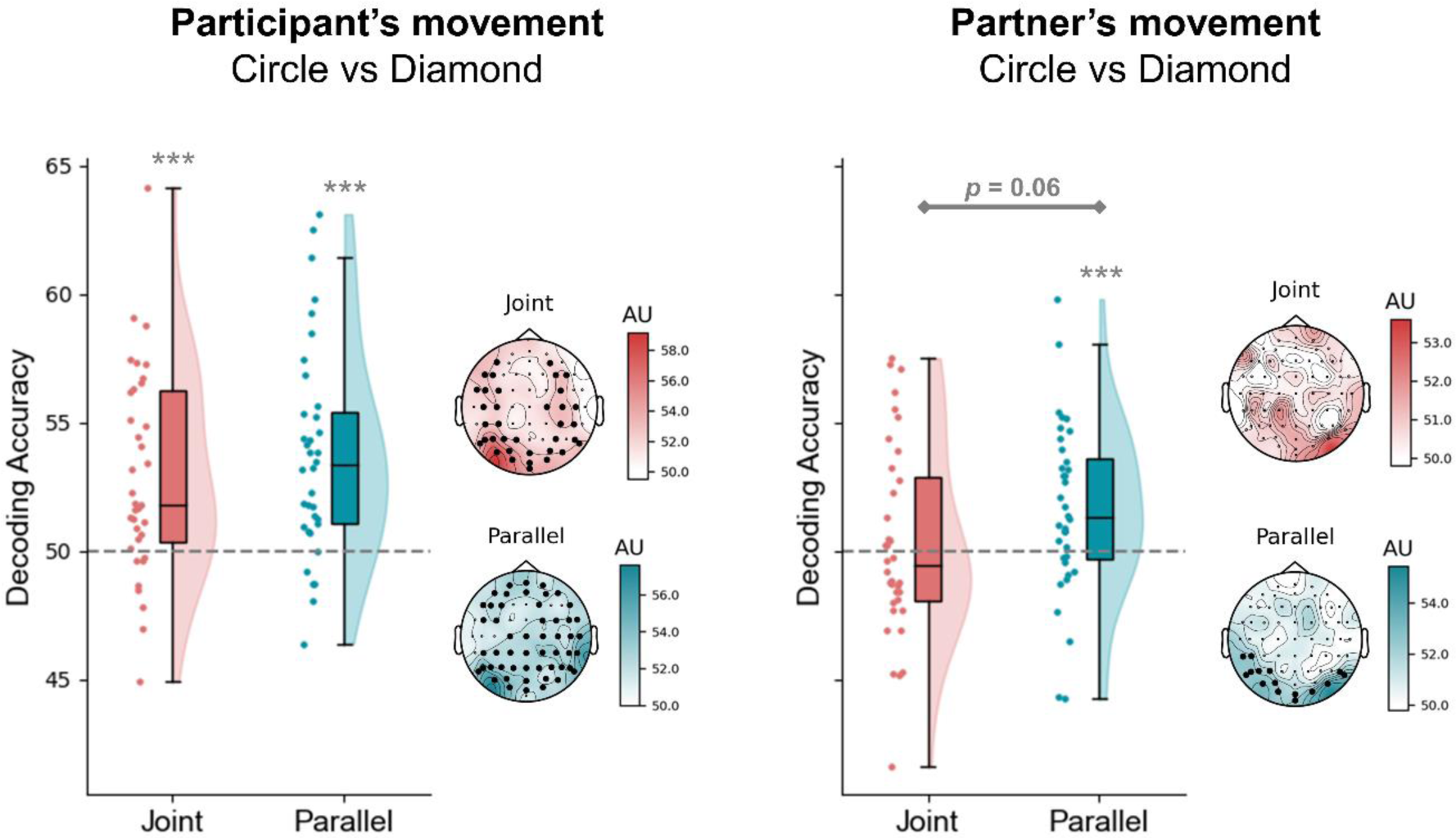
Decoding of Participant’s and Partner’s Movements. *Note. Left panel*: accuracies resulting from the decoding of the instructed participant’s movement (Circle or Diamond), separately for Joint and Parallel task. *Right panel*: accuracies resulting from the decoding of the instructed partner’s movement (Circle or Diamond), separately for Joint and Parallel task. See Figure 5 for plotting conventions.

The prepared participant’s movement was successfully classified above chance in both the Joint (t_35_ = 4.44, *p* < 0.01, d = 0.74, BF_10_ > 100) and Parallel task (t_35_ = 5.54, *p* < 0.01, d = 0.92, BF_10_ > 100). We found no evidence indicating that the strength of the participant’s movement representation differed between Joint and Parallel contexts (t_35_ = 0.79, *p* = 0.431, d = 0.20, BF_10_ = 0.24). Spatially, clusters of above-chance electrodes were widespread over the whole scalp.

Interestingly, classifying the partner’s movement lead to above chance accuracies in the Parallel task (t_35_ = 2.92, *p* < 0.01, d = 0.49, BF_10_ = 12.92), but not in the Joint task (t_35_ = 0.44, *p* = 0.33, d = 0.07, BF_10_ = 0.40). Decoding accuracies of the partner’s movement were almost significantly larger in the Parallel compared to the Joint task, although the formal significance threshold was not met (t_35_ = 1.90, *p* = 0.06, d = 0.38, BF_10_ = 0.90). Consistently, the spatial decoding revealed an occipital cluster of electrodes significantly above chance only in the Parallel task.

## Discussion

According to recent proposals, engaging in successful coordinated interactions entails the proactive anticipation of our partner’s actions (Kilner et al., 2007), and their integration with our own motor contribution in a Dyadic Motor Plan towards the shared goal (Sacheli et al., 2018). This perspective is in line with the idea of engaging in a ‘we-mode’ during social interactions (Gallotti & Frith, 2013), intended as a task representation transcending the individual parts and encompassing both agents. However, so far the emergence of such integrated representations in preparation for Joint Actions has not been tested directly, but rather was supported by reduced interference effects on overt behavior (Clarke et al., 2019; Rocca et al., 2023; Sacheli et al., 2018). At the neural level, previous studies did not address specifically the representational content of this hypothesized anticipatory activity. EEG studies targeting specifically the preparatory phase preceding Joint Actions reported only univariate differences attributed to the interactive task context (Kourtis et al., 2013, 2014, 2019), remaining agnostic with respect to the informational content of such neural activity.

In contrast, ours is the first study of Joint Actions in humans shifting the focus from the execution and observation phase to the preparation phase, and using multivariate analyses to directly probe the representational content of the electrophysiological activity before the movements take place. We reasoned that an intertwined encoding of the participant’s and the partner’s prospective contributions into a Dyadic Motor Plan would result in their lower discriminability. Confirming this hypothesis, the multivariate approach we implemented revealed that the two incongruent movement combinations (i.e., circle-diamond and diamond-circle) were significantly easier to discriminate when the two agents were acting in parallel, compared to jointly. This finding provides compelling evidence for the formation of an integrated representation incorporating both upcoming movements. In turn, and consistently with previous studies, this lead to reduced VMI, quantified in the distortion of the drawn trajectories, as the active anticipation of observing an incongruent movement ensures its diminished interfering effect (Kaiser & Schütz-Bosbach, 2018).

We further tested to what extent each of the two individual contributions was decodable from the pattern of neural activity recorded during the preparation interval. The upcoming movement of the participant was successfully decodable in both social contexts, indicating that the motor plan associated with the shape to draw was being strongly prepared (Ariani et al., 2015, 2022). With respect to the action of the partner, in the Joint condition the classifier failed at discriminating whether they were about to draw a circle or a diamond. On the contrary, this same contrast yielded high decoding accuracies during Parallel interactions, indicating that in this condition the upcoming partner’s movement was represented in a format that could be more effectively picked up by the classifier. Crucially, good performance in catch trials detection clearly showed that participants were efficiently maintaining the information pertaining the movements of the partner. This excluded the possibility of our results being explained by an attentional mechanism. Namely, low classification accuracies in the joint social context cannot be due to the action information from the partner being entirely disregarded. Moreover, we consider it highly implausible that participants directed more attention towards the partner’s action contribution in the parallel condition, as the task did not require to adjust their behavior based on the partner’s. We think these results represent an important step forward in understanding how the available information on the partner’s movement is represented as a function of the interactive social context. Previous studies focused predominantly on motor activation in response to an observed movement or during interaction, often in situations in which participants had no prior information concerning the partner’s contribution (Bolt & Loehr, 2021). As a notable example, Sacheli and colleagues (2019) revealed a stronger hemodynamic response in the left ventral premotor cortex, tracking the identity of the unfolding partner’s movement, while agents took turns in playing a melody (Sacheli et al., 2019). Conversely, we investigated the anticipatory phase of Joint Actions, and provided complementary evidence showing that the upcoming movement of an interactive partner could not be successfully decoded in the interval preceding its unfolding. This finding is consistent with the idea that, when cued in advance, the other’s contribution is anticipated integrated to one’s own action and in reference to it, a representational format that does not allow for its classification in isolation but is functional to minimize its subsequent interfering effect. Therefore, our results extend on previous research in suggesting that motor interactions rely on complex representational dynamics, potentially undergoing reorganization from the anticipation to the implementation phase (Ariani et al., 2022; Elsayed et al., 2016), and shaped by social context, information availability, and its functional relevance.

With respect to the spatial distribution of our results, the decoding analysis suggests that occipital electrodes are those carrying more information concerning the partner’s upcoming action in the Parallel condition. This topography hints at a larger recruitment of areas associated with visual processing to maintain the partner’s contribution throughout the delay period (Christophel et al., 2017). One possible explanation for this might be that in the Parallel condition participants are mostly anticipating the expected visual outcome of the partner’s action, rather than its kinematic properties. On the other hand, there is growing evidence showing that the maintenance in working memory (WM) of information concerning motion patterns, biological movements, and hand postures elicits activation of somatosensory cortices, as indexed by suppression of rolandic mu rhythm (Christophel & Haynes, 2014; Galvez-Pol et al., 2018; Gao et al., 2015). Moreover, the role of motor activity to smoothly coordinate with an interaction partner has been widely emphasized by previous studies on Joint Action (Bolt & Loehr, 2021). In our data, we did not observe larger decoding accuracies in the joint condition over sensorimotor cortices, nor in any other scalp location. Nevertheless, we speculatively raise the intriguing hypothesis that the social context affects the representational format of the information concerning the partner’s movement. Research in the field of WM is strongly demonstrating how task demands influence the encoding and maintenance of items, supporting the idea that information is coded in the most suitable way to effectively drive behavior (Henderson et al., 2022; Myers et al., 2017; van Ede & Nobre, 2023). In keeping with the premises of the Dyadic Motor Plan, we claim that in an interactive context the optimal representational format for the partner’s action is motoric in nature, already during the anticipation phase. This is functional to allow its efficient assimilation with the participant’s contribution. In fact, perhaps unsurprisingly, the results on the decoding of the participant’s own upcoming movement are compatible with a more motoric representational format, showing widespread scalp areas of high decoding accuracies in both social contexts, including also anterior electrodes over sensorimotor cortices. On the contrary, acting alongside does not require the motoric encoding of the other’s action, as this does not need to be integrated with the motor plan of the participant, and can be maintained for future recognition as more visual or abstract semantic content.

In conclusion, our current results support the Dyadic Motor Plan framework and extend on it, demonstrating that the neural representations of the upcoming joint configuration and of the partner’s movement are affected by the social context manipulation. These findings suggest that engaging in Joint Action induces the reformatting of the encoded information into integrated motoric representations, to optimize the subsequent coordinated behavior. Disentangling the specific formats of the encoded actions, and investigating how they evolve between preparation and implementation, thus remain open and fascinating avenues for future research.

## Supplementary Materials

### Control Analysis on VMI and Drawing Time

We fitted an additional model including the centered factor Drawing Time and its interactions with the factors Social context and Congruency. The rationale of fitting this model was accounting for a potential effect of the speed of drawing on the distortion in the trajectories. While the effects of Social context, Congruency and their interaction were unaffected by the inclusion of the factor Drawing time, we found a significant interaction of Congruency and Drawing Time (β = -0.78, 95% CI =-1.27, -0.30], t_16625_ = -3.18, *p* = 0.001). This interaction indicates that slower movements are associated with larger distortions in incongruent trials, discarding the possibility of a speed-accuracy trade-off between movement duration and variability in drawn trajectories.

**Figure S1.**
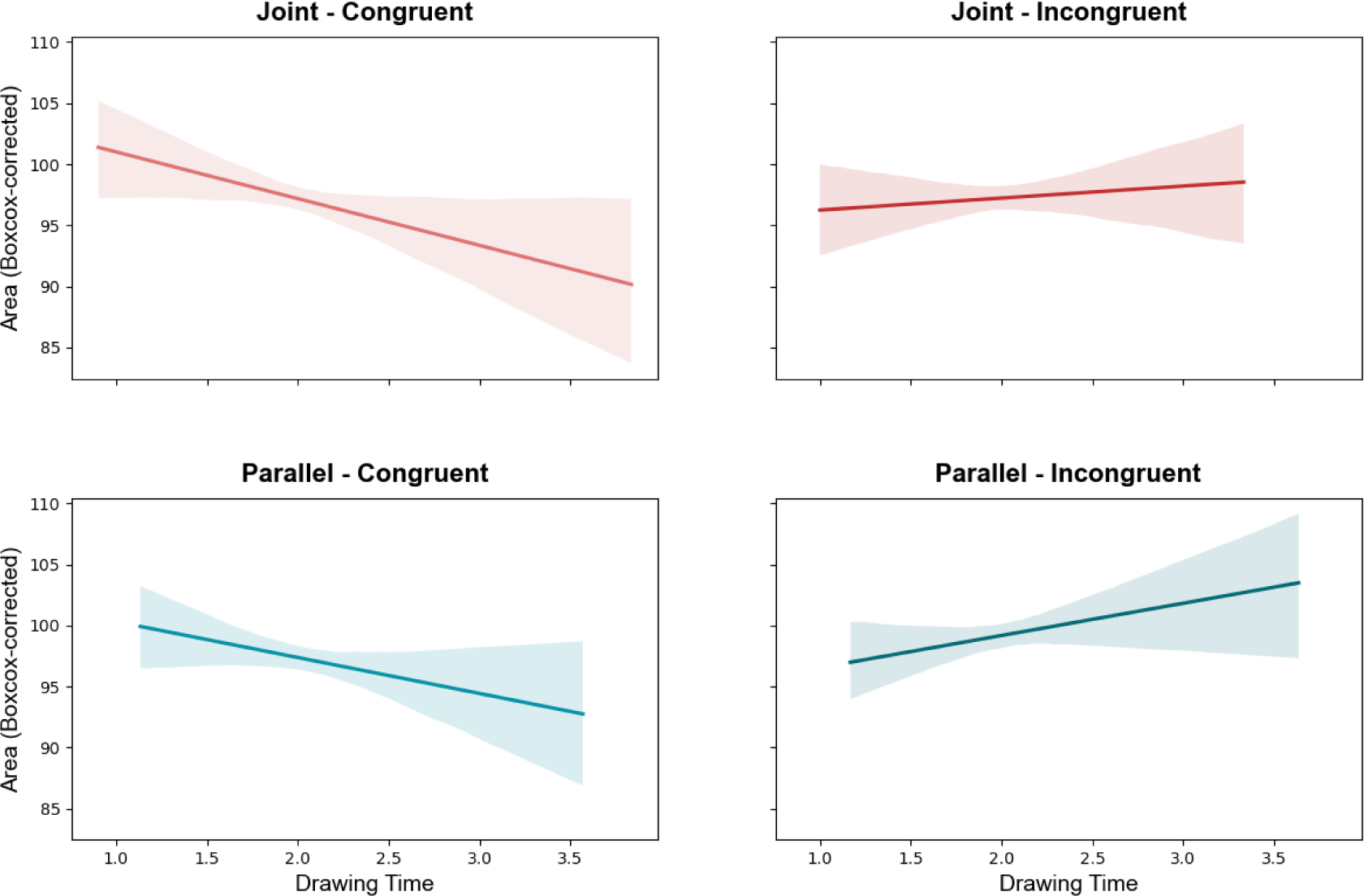
Control Analysis on VMI and Drawing Time. *Note*. The significant interaction of the factor Congruency and Drawing Time suggests that larger VMI is observed in incongruent trials with longer Drawing Time, as opposed to congruent trials. This indicates that a phenomenon akin to ‘speed-accuracy trade-off’ (i.e., poorer performance in favor of faster execution) can be observed only in congruent but not in incongruent trials.

## Significance statement

Coordinating our actions with that of other agents is a common yet rather complex endeavor. It has been proposed that, to engage in successful Joint Actions, we proactively anticipate the action plans of our partner, which become integrated to our own. As a consequence, the other’s movement interferes less with the execution of our own movements. In the present study, we demonstrate that, depending on whether the social context is interactive or not, the information on each agent’s action plan is encoded differently in the pattern of brain activity preceding the action. Our findings support the hypothesis that smooth coordination in social settings relies on predictive and integrative processes.

## Acknowledgements

SF was supported by the German Research Foundation (Deutsche Forschungsgemeinschaft, DFG) under Germany’s Excellence Strategy-EXC 2002/1, Science of Intelligence (Project Ref.: 390523135) and the Einstein Foundation Berlin. MB is supported by an Einstein Strategic Professorship (Einstein Foundation Berlin).

## Data and Code availability statement

Raw data and analyses scripts will be made publicly available upon publication through the Open Science Framework

## Authors contributions

**Silvia Formica**: Conceptualization, Investigation, Analyses, Writing. **Marcel Brass**: Conceptualization, Supervision, Writing.

